# Optimal sigmoid function models for analysis of transspinal evoked potential recruitment curves recorded from different muscles

**DOI:** 10.1101/2024.07.15.603573

**Authors:** Andreas Skiadopoulos, Maria Knikou

**Author notes:** Corresponding Author (AS).

## Abstract

Recruitment input-output curves of transspinal evoked potentials that represent the net spinal motor output of alpha motor neurons, and those of cortically, spinally or peripherally induced responses are used extensively in research as neurophysiological biomarkers to establish physiological or pathological behavior of neuronal networks as well as post-treatment recovery. A comparison between different sigmoidal models to fit the transspinal evoked potentials recruitment curve and estimate the parameters of physiological importance has not been performed. This study sought to address this gap by fitting eight sigmoidal models (Boltzmann, Hill, Log-Logistic, Log-Normal, Weibull-1, Weibull-2, Gompertz, Extreme Value Function) to the transspinal evoked potentials recruitment curves of soleus and tibialis anterior recorded under four different cathodal stimulation settings. The sigmoidal models were ranked based on the Akaike information criterion, and their performance was assessed in terms of Akaike differences and weights values. Additionally, an interclass correlation coefficient between the predicted parameters derived from the best models fitted to the recruitment curves was also established. A Bland-Altman analysis was conducted to evaluate the agreement between the predicted parameters from the best models. The findings revealed a muscle dependency, with the Boltzmann and Hill models identified as the best fits for the soleus, while the Extreme Value Function and Boltzmann models were optimal for the tibialis anterior transspinal evoked potentials recruitment curves. Excellent agreement for the upper asymptote, slope, and inflection point parameters was found between Boltzmann and Hill models for the soleus, and for slope and inflection point parameters between Extreme Value Function and Boltzmann models for the tibialis anterior. Notably, the Boltzmann model for soleus and the Extreme Value Function model for tibialis anterior exhibited less susceptibility to inaccuracies in estimated parameters. Based on these findings, we suggest the Boltzmann and the Extreme Value Function models for fitting the soleus and the tibialis anterior transspinal evoked potentials recruitment curve, respectively.

## Introduction

The input-output relationship between the stimulation intensity and response amplitude has a sigmoidal (or S) shape for the corticospinal motor, spinally mediated monosynaptic reflex, and peripheral motor pathways. Specifically, the motor evoked potentials (MEPs), H-reflex (from threshold up to maximal amplitudes), transspinal evoked potentials (TEPs) recorded synchronously from flexor and extensor knee and ankle muscles, and peripheral motor M-waves all exhibit a sigmoidal input-output relation [1–10]. Although for each response different neuronal pathways are engaged during depolarization of spinal alpha motoneurons, all four types of compound muscle action potentials manifest in a nonlinear input-output relation.

The sigmoid or S-shaped curve is the result of a combination of factors including but not limited to the well-established physiological principle that stimuli at increasing strength recruit motoneurons with increasing motor unit potentials [5,11]. Afferent volleys initiated in the muscle-nerve junction and responses recorded in ventral roots have a clear S-shape [12]. Additionally, the exponential distribution of recruitment thresholds across the motoneuron pool [13], discharge probability of multiple motoneurons/interneurons [2], intrinsic threshold of motoneurons [14], input resistance [15,16], relative share of synaptic inputs whereas a motoneuron that receives a larger synaptic input tends to be recruited before cells with smaller inputs [17], amplification of synaptic input by persistent inward currents [17], excitability properties of afferents, and discharge probability of single motor units [18] all contribute to the S-shaped input-output relation.

Nonlinear and linear regression models are used interchangeably to predict parameters of physiological importance that aid to understand the function of the nervous system in health and disease [6,19–25]. These parameters include the stimulation intensities corresponding to threshold, 50 % and 100 % of MEP, H-reflex, TEP, and M-wave amplitude, as well as the slope (known as gain) of the curve at the inflection point at x-axis [26–28]. The best fitted models for the soleus (SOL) H-reflex and M-wave input-output curves are the cubic splines, ninth-order polynomial, Boltzmann (logistic) and logarithmic functions, as determined by goodness-of-fit statistics (r-squared, root-mean-squared error), when compared to linear and power fit models [3]. The Boltzmann model is the most robust and easy to interpret because it estimates parameters that are physiologically meaningful [3]. The TEP input-output curve has been modeled with both Boltzmann and Hill functions [8,19,21,29] and a sixth-order polynomial function [22,23]. However, there is currently no evidence on the most optimal mathematical model to fit the TEP input-output curves.

Use of nonlinear regression models has significant advantages. Nonlinear models are more realistic compared to linear and polynomial models and can effectively capture the characteristics of input-output curves using only a few key parameters. Linear models are unable to account for saturation or plateau in the S-shaped recruitment curves, and high-order polynomial models require larger number of parameters compared to a three-parameter logistic model to achieve similar fitted values. Additionally, polynomial functions may produce unreliable predictions or interpolations that could result in negative responses [22]. Thus, linear, and polynomial functions were excluded from this study.

Taken altogether, in this study we fitted the TEP recruitment curves with eight different sigmoidal models. For the 3 out of the 8 models the rate of change was symmetric about their inflection point (Boltzmann, Hill, and Log-Logistic models), and asymmetric for the remaining models (Gompertz, Extreme Value Function (EVF), Weibull-1, Weibull-2 and Log-Normal). Nonlinear models were used to provide a more parsimonious description of the data compared to polynomials (i.e., better data fit and more precise parameters), and estimate parameters that have physiological meaning [30]. The objective of this study was to identify the nonlinear model that best fits the TEP recruitment input-output curves based on information-theoretic criteria [30–33]. The nonlinear models were fitted to the right and left SOL and tibialis anterior (TA) TEP input-output curves assembled under four different positions and number of the cathodal electrode for transspinal stimulation [6].

## Material and Methods

### Experimental dataset

In this study, a total of eight nonlinear models were fitted to the left and right SOL and TA TEP recruitment input-output curves previously assembled in a cohort (6 females; 5 males; age 25.7 ± 5.2 years; height 1.7 ± 0.1 m; mass 70.2 ± 12.1 kg) of heathy subjects [6]. The recruitment input-output curves were assembled by evoking four TEPs at each increasing stimulation intensities in steps of 0.5 mA, starting from below the right SOL TEP threshold until either the TEP amplitude reached a plateau or transspinal stimulation produced discomfort. With subject’s supine, the TEP input-output curves were assembled when the transspinal cathode electrode was one rectangular electrode placed at midline (Protocol Knikou-Lab4Recovery; P1), one square electrode placed at midline (P2), two square electrodes with 1 cm apart placed at midline (P3), and one square electrode placed on each paravertebral side (P4). The TEP amplitude was measured as the area under the full wave rectified action potential expressed in mV × ms (Fig. 1). The experimental protocol was approved by the City University of New York (New York, NY) Institutional Review Board Committee (IRB No. 2019-0806) and was conducted in compliance with the Declaration of Helsinki. All participants provided written informed consent before participation in the study. The experimental procedure for transspinal stimulation is described in detail in [6].

**Fig 1.**
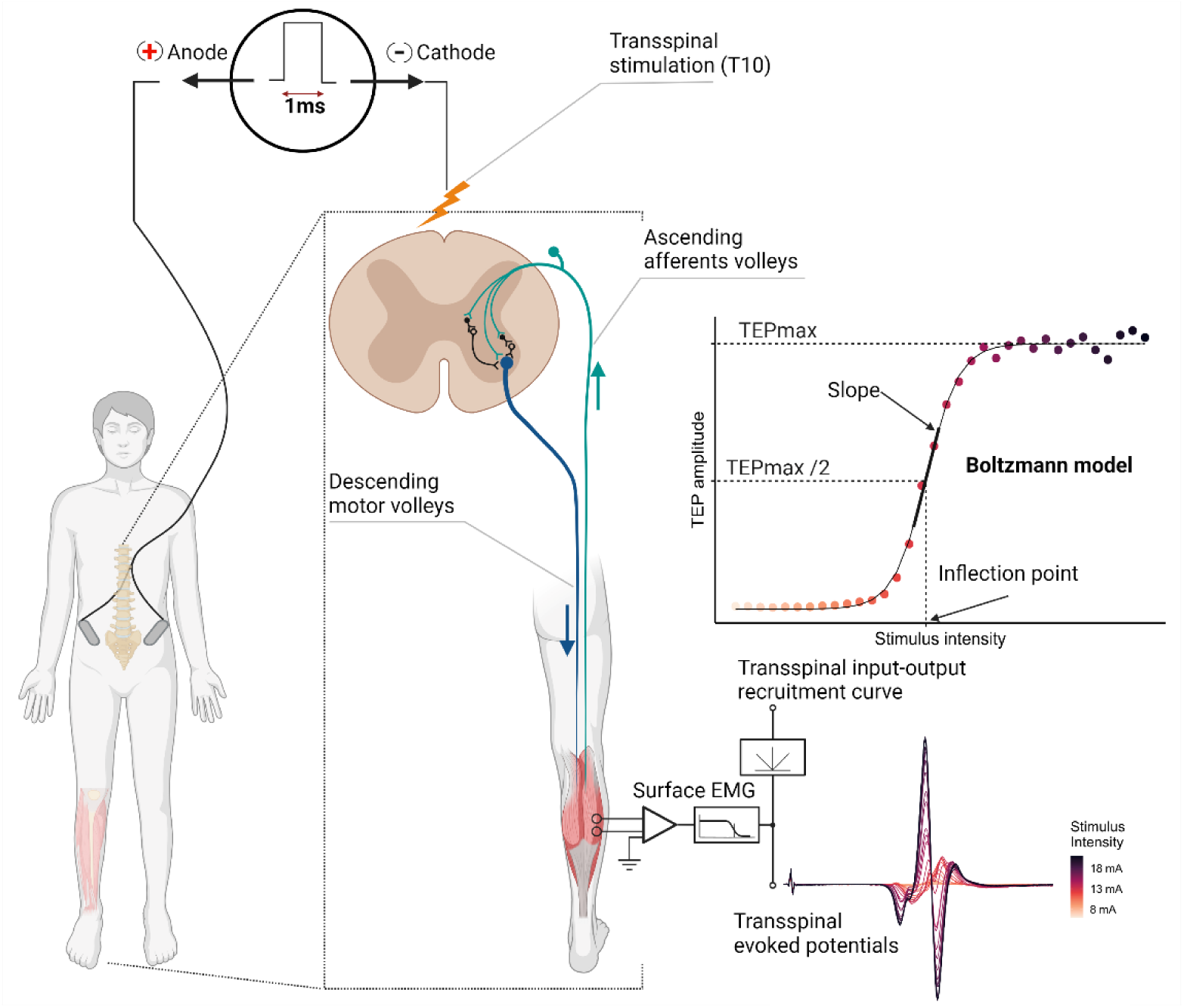
Schematic Illustration of Transspinal Stimulation. Example of recordings and recruitment of transspinal evoked potentials (TEPs) when delivering a single pulse of transspinal stimulation with increasing intensity. To quantify each muscle’s transspinal input-output recruitment curve, the TEP amplitude (area under curve) was plotted as a function of the stimulus intensity and a nonlinear regression model was fitted to the data (created with BioRender.com).

### Nonlinear mathematical models for TEP input-output curves

The eight nonlinear models used to establish the best-of-fit for the right and left SOL and TA TEP input-output curves are indicated in Table 1. All models follow a sigmoidal shape and have an upper asymptote corresponding to the maximum TEP amplitude (TEPmax). The point of maximum steepness occurs at the inflection point of x-axis and represents the recruitment gain. The inflection point is the point on the X-axis (abscissa) at which the first derivative of the input-output curve is maximized. The inflection point shows at which stimulation intensity the increasing growth rate of TEP amplitude turns to a decreasing rate. A difference between the models is that some exhibit symmetry on each side of the inflection point. This characteristic makes symmetrical models less flexible compared to asymmetrical sigmoid models, potentially limiting their capacity to accurately describe the TEP input-output recruitment curves.

**Table 1.**
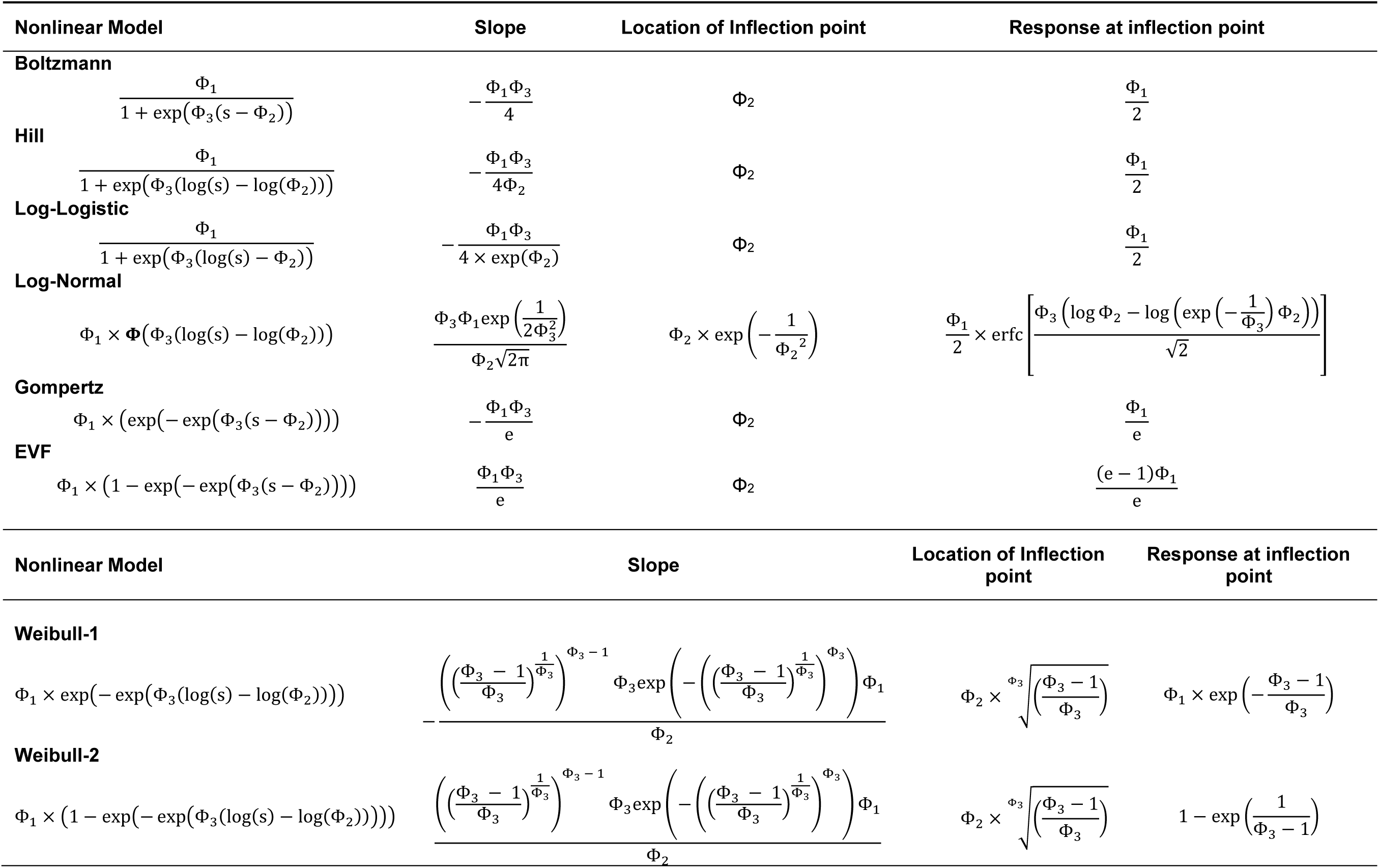
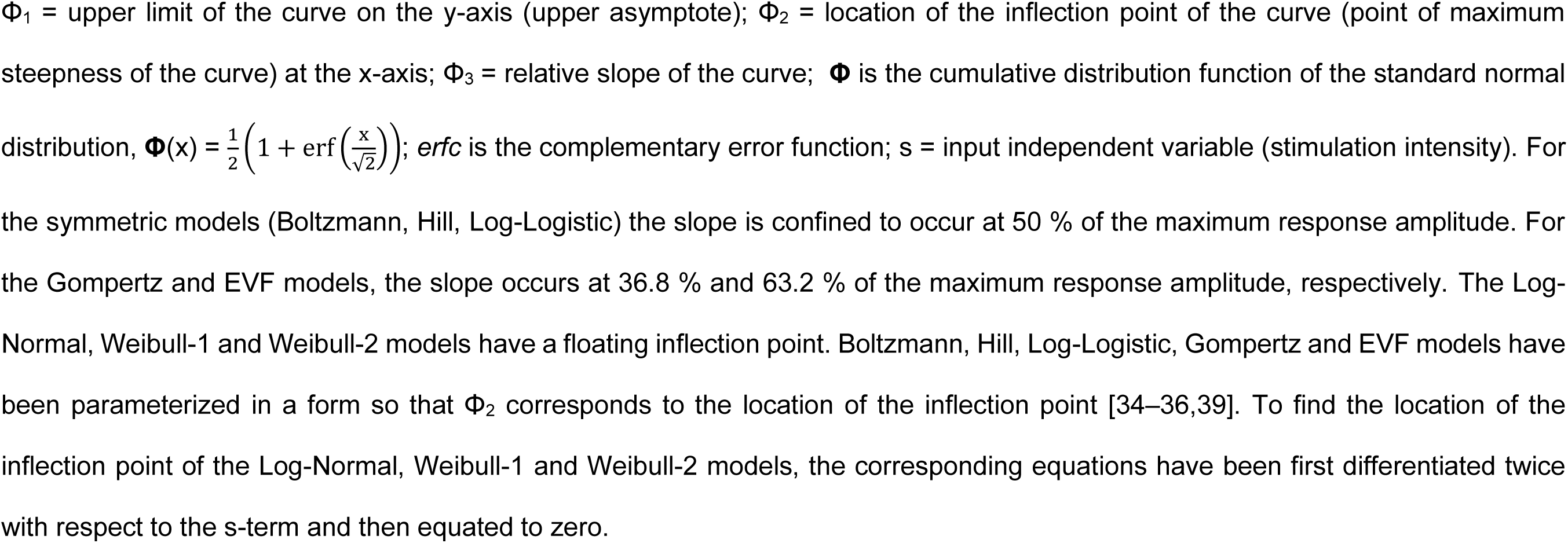
Modelling TEP recruitment curves: equations and derived parameters.

The Boltzmann, Hill and the Log-Logistic models are symmetrical and exhibit a fixed inflection point at 50 % of the maximum TEP amplitude [34,35]. The inflection point for these models can be interpreted as the stimulation intensity at which half of the maximum TEP amplitude is achieved. In contrast, the Gompertz and EVF models are asymmetrical about the inflection point. The inflection points of the Gompertz and EVF models are fixed and correspond to stimulation intensities at which the TEP amplitude increases by 36.8 % and 63.2 % of the maximum TEP amplitude, respectively [36,37]. The Log-Normal, Weibull-1 and Weibull-2 models are more flexible compared to the other models because they have a floating inflection point, and its location varies with the relative slope parameter [34,35,38]. The EVF and Weibull-2 models are well-suited for describing recruitment curves with a slower rise of slope reaching a maximum, followed by a more rapid decline. In contrast, the Weibull-1 and Gompertz models exhibit a quick initial rise of the slope followed by a gradual decline [35].

### Nonlinear regression analysis

For all nonlinear regression models, a three-parameter function that corresponds to the TEP amplitude, y, at stimulation intensity, s, which can be generally given by the mean function, *f*, was defined as indicated in Eq 1,

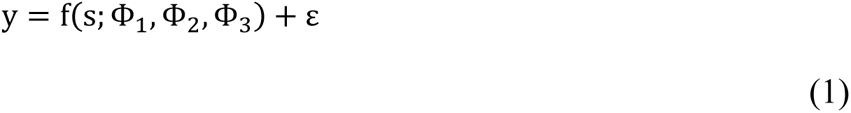

where the additive error, denoted with ε, is assumed to be an uncorrelated stationary process with zero mean and constant variance σ^2^. In Eq 1, the Φ_1_ parameter denotes the upper limit or maximum horizontal asymptote, which reflects the TEP amplitude value at saturation (TEPmax), while the Φ_3_ parameter is the relative slope around the inflection point, Φ_2_. The nonlinear regression models were fitted to the stimulation-response data by nonlinear least-squares method using the *minpack.lm* and *drm* R packages. The objective was to determine the model that best describes the systematic TEP input-output trend. To address this objective, separate nonlinear models were fitted to each single-subject data. The nonlinear models were fitted to the sample means obtained from averaging the individual pseudo-replicates of TEP responses at each stimulation intensity. Whether fitting the nonlinear regression models to mean values or to individual pseudo-replicates, the same parameters are obtained (i.e., same curves), assuming same number of pseudo-replicates at each stimulation intensity. The assumptions of the models were evaluated using graphical procedures [40] and formal statistical tests. Variance homoskedasticity was assessed by examining plots of standardized residuals against fitted values. The Shapiro-Wilk and the Ljung-Box test were used to evaluate normality and residual correlation (first-order autoregressive correlation).

### Model selection based on information-theoretic criteria

The information-theoretic approach was used to select the most optimal model in terms of Kullback-Leibler information for describing the TEP recruitment curve [30,31]. The second order Akaike information criterion (AICc) was used because is suitable for small sample sizes without extremely nonnormal distribution [30]. The models were ranked based on the AICc value [41], and rated according to their Δ_AICc_ values and Akaike weights (W_AICc_). The Δ_AICc_ values were calculated as the AICc difference between a specific model and the model with the lowest AICc value. Using the Δ_AICc_ differences the relative likelihood of each model was determined, which was then normalized to calculate the Akaike weights [30,31]. The Akaike weights are interpreted as the probability of each model being the most optimal, based on the available data and models. The evidence ratio of Akaike weights was computed to quantify the amount of support of the best model M_y_ relative to another model M_x_ (evidence ratio = W_AICc_ of M_y_ / W_AICc_ of M_x_). A 95 % confidence set of models that were most likely the best approximating models was established by summing the Akaike weights from largest to smallest until the sum was ≥ 0.95 (Burnham et al., 2002). The best AICc-selected models were further analyzed using the Jackknife leave-one-out resampling method, and the corresponding Akaike’s weights (JK W_AICc_) values were calculated. Goodness-of-fit was also evaluated using the pseudo coefficient of determination (R^2^) averaged by use of Fisher’s Z-transformation [42].

The most optimal mathematical models were further considered based on their ability to provide reliable estimates of the physiological parameters (TEPmax, slope, and inflection point) of TEP recruitment input-output curves. Two-way mixed effects interclass correlation coefficient was used to determine the degree of agreement (ρ) between the best models (degree of agreement was poor for ρ < 0.5, moderate for 0.5 ≤ ρ < 0.75, good for 0.75 ≤ ρ ≤ 0.9, and excellent for ρ > 0.9) [43]. The degree of agreement between estimates of parameters obtained by two different models was further evaluated with Bland-Altman analysis by expressing paired differences as percentages of their mean value [44]. TEP threshold was defined as the minimum stimulation intensity required to recruit the most excitable motor neuronal structure [45]. All analyses were conducted in R 4.2.3 [46].

## Results

In Fig 2, the right and left TA and SOL TEP recruitment input-output curves assembled for each subject and stimulation protocol are shown. It is apparent that the S-shape is shifted more to the right (i.e., horizontal shifting of the TEP recruitment curve) for some subjects and protocols. These results are consistent with subject dependency of excitation threshold and recently shown a shift to the right from P1 to P4 [6].

**Fig 2.**
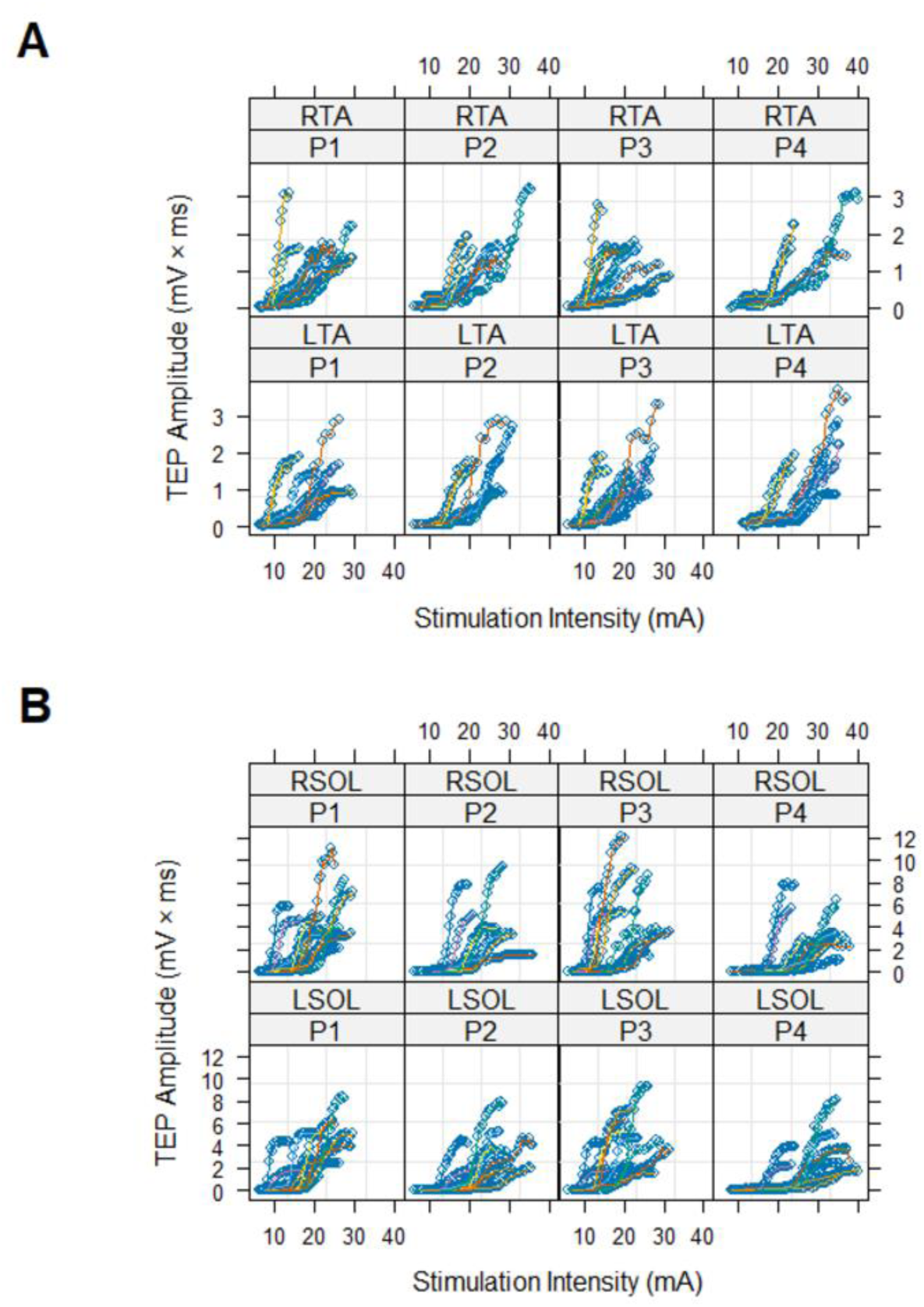
TEP recruitment curves. Right and left TA (A) and SOL (B) transspinal evoked potentials (TEPs) input-output curves assembled for each subject and stimulation protocol. Points corresponding to data from the same subject are connected by lines. Within- and between-subject variability in stimulation intensities required to reach the same level of TEP amplitude, TEP threshold, maximum TEP amplitude, and the intensity required to reach maximal amplitudes are evident. RTA = right tibialis anterior; LTA = left tibialis anterior; RSOL = right soleus; LSOL = left soleus; P1, P2, P3, P4 = stimulation protocols.

In Fig 3, a representative TEP recruitment input-output curve assembled for the right SOL muscle under protocol P3 is shown for a single subject. The TEP input-output recruitment curve is shown while fitted with the Boltzmann, Hill, Log-Logistic, Log-Normal, Weibull-1, Weibull-2, EVF, and Gompertz nonlinear regression models. For all cases the AICc and R^2^ values are also shown. The filled circle in each graph corresponds to the model’s slope estimated using the equations presented in Table 1. The sigmoidal shape of the TEP input-output curve identified by all eight nonlinear regression models is further supported by the bell-shape curve, shown with dotted lines, of the amount-of-change function (first derivative of the TEP recruitment input-output curve). The amount-of-change curves were calculated as the finite difference of models’ fitted data [47]. The amount-of-change curve is symmetrical on each side of the inflection point for the Boltzmann, Hill, and the Log-Logistic models. The amount-of-change curve demonstrated positive skewness for the Weibull-1 and the Gompertz asymmetrical models, whereas the Weibull-2 and the EVF models exhibited negative skewness. The Log-Normal model, despite its asymmetry, exhibits a similar shape to that observed in symmetrical models.

**Fig 3.**
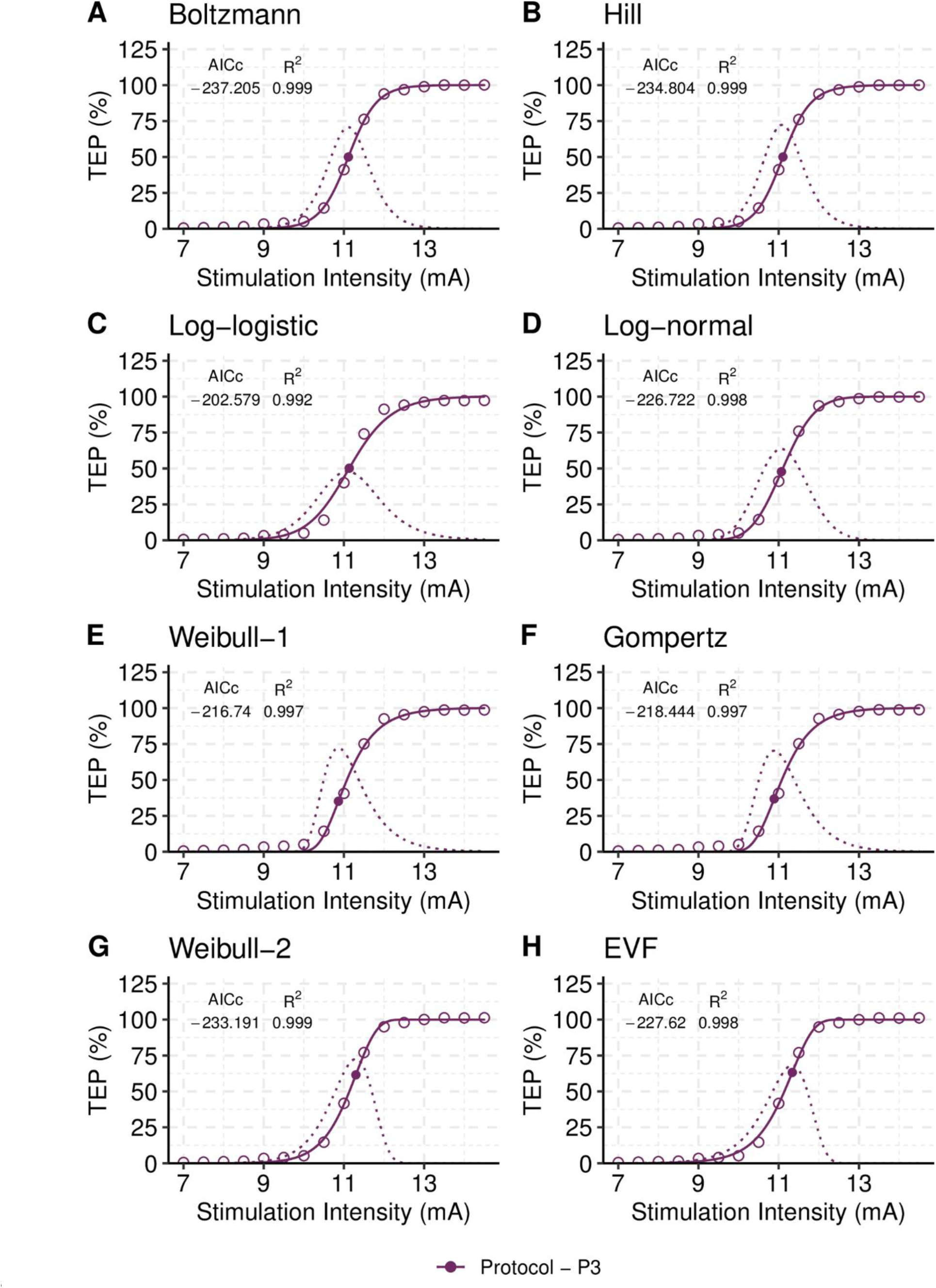
Modelling the TEP recruitment curve of a representative single subject. Each plot (A-H) contains the stimulation-response data (circles), the predictions obtained from the fitted models, and the amount-of-change curve (bell-shaped curve). The stimulation intensity corresponding to the peak of the amount-of-change curve corresponds to that of the inflection point of the fitted model (showed with a filled circle on each curve). AICc and R^2^ values are presented alongside the plots to assess goodness-of-fit. The Boltzmann model is the best model in terms of Kullback-Leibler discrepancy (AICc = -237.205). The R^2^ value was found to be the same for all models, rendering it unreliable for model selection. To ensure comparability, the fitted curves were rescaled to the maximum predicted value of each model, ranging from 0 % to 100%. The data is for transspinal evoked potentials (TEPs) of the right SOL from a single subject under protocol P3.

Table 2 presents the ranking of the models fitted to the single-subject TEP input-output curves shown in Fig 3. The best model in terms of Kullback-Leibler discrepancy is the Boltzmann because it had the lowest AICc value of all models (AICc = -237.205). The inflection point, or S50, of the Boltzmann model corresponds to stimulation intensities of 11.11 ± 0.02 mA (mean ± standard error) that results in TEP responses of 3.75 ± 0.05 mV × ms (estimated using equations in Table 1). Note that the inflection points of the symmetrical models, which include Boltzmann and Hill models, are located at stimulation intensity of 50 % of TEPmax (thus denoted as S50). The Boltzmann slope at the inflection point is 5.27 ± 0.22 mV × ms / mA. The Akaike’s weight for the Boltzmann model (W_AICc_ = 0.69) was the largest relative to the Akaike’s weights for the other ranked models (e.g., the Akaike’s weight of Hill model is W_AICc_ = 0.21). This indicates that there is a 69 % chance that the Boltzmann model is the best approximating model for describing the given data, while the Hill model has 21 % chance and Weibull-2 model has only 9 % chance.

**Table 2.** Evaluation of models and parameters for describing the TEP recruitment curve of a representative single subject.

| Model | AICc | $\Delta_{AICc}$ | $L_{AICc}$ | $W_{AICc}$ | Evidence | TEPmax<sup>†</sup> | Slope<sup>‡</sup> | S50<sup>♀</sup> |
| --- | --- | --- | --- | --- | --- | --- | --- | --- |
| Log-logistic | -202.579 | 34.63 | 0.00 | 0.00 | $33 \times 10^6$ | 7.61 (0.17) | 3.67 (0.32) | 11.12 (1.00) |
| Weibull-1 | -216.740 | 20.46 | 0.00 | 0.00 | 27,780.7 | 7.58 (0.11) | 5.45 (0.41) | 11.08 (0.03) |
| Gompertz | -218.444 | 18.76 | 0.00 | 0.00 | 11,852.2 | 7.55 (0.10) | 5.31 (0.39) | 11.08 (0.34) |
| Log-normal | -226.722 | 10.48 | 0.01 | 0.00 | 188.9 | 7.38 (0.07) | 4.75 (0.25) | 11.14 (0.02) |
| EVF | -227.620 | 9.59 | 0.01 | 0.01 | 120.6 | 7.44 (0.07) | 5.03 (0.25) | 11.10 (0.03) |
| Weibull-2 | -233.191 | 4.01 | 0.13 | 0.09 | 7.4 | 7.37 (0.06) | 5.35 (0.25) | 11.14 (0.02) |
| Hill | -234.804 | 2.4 | 0.30 | 0.21 | 3.3 | 7.46 (0.06) | 5.39 (0.25) | 11.10 (0.02) |
| Boltzmann | -237.205 | 0.0 | 1.00 | 0.69 | 1.0 | 7.45 (0.05) | 5.27 (0.22) | 11.11 (0.02) |

AICc = second order Akaike Information Criterion; $\Delta_{AICc}$ = AICc - min(AICc); $L_{AICc}$ = relative likelihood for model; $W_{AICc}$ = Akaike weight (i.e., normalized $L_{AICc}$ ); Evidence = max( $W_{AICc}$ ) / $W_{AICc}$ . $\Delta_{AICc}$ represents the difference between a particular model and the model with the lowest AICc value, indicating the amount of information lost when using the specific model. $W_{AICc}$ represents the probability of a given model being the best model over the model with the lowest AICc score. Evidence ratio refers to the relative likelihood of model pairs and provides information about the fitted models in terms of which one is better in terms of Kullback-Leibler information.
<sup>†</sup> Units: mV × ms
<sup>‡</sup> Units: mV × ms / mA
<sup>♀</sup> Units: mA

The S50 of the second-best Hill model corresponds to stimulation intensity of 11.10 ± 0.02 mA that results in TEP response of 3.70 ± 0.10 mV × ms, which are similar to those obtained from the Boltzmann model. The inflection points of the third-best model, Weibull-2, corresponds to stimulation intensity of 11.30 ± 0.02 mA, which results in a TEP response of 4.50 ± 0.10 mV × ms (note that Table 2 shows values only at S50 for comparison). Weibull-2 is not a symmetrical model and S50 is a point different than the inflection point. The S50 value for the Weibull-2 model is 11.14 ± 0.02 mA (Table 2). The best three models estimate comparable TEP responses at S50 – i.e., stimulation intensity at 50 % of TEPmax. The slope at S50 for the Hill and Weibull-2 models is 5.39 ± 0.25 and 5.35 ± 0.25 mV × ms / mA, respectively (Table 2). The other models represent poor approximations to the given data (i.e., the models in Table 2 with Δ_AICc_ ≥ 8.16). Although all fitted models satisfied the goodness-of-fit criterion of coefficient of determination (R^2^ > 0.996), the AIC-based model selection approach revealed that the Boltzmann model is the best model for the representative SOL TEP input-output recruitment curve from a single subject shown in Fig 3.

### Model selection for TEP recruitment input-output curves

In Table 3 the ranking of the models based on the average of the AICc values across subjects is shown. Both Boltzmann and Hill models were the two-best models for the right and left SOL TEP recruitment input-output curves for all four protocols. The evidence ratio between the two best models ranged from 1.3 to 3.0, indicating similar support for both models (i.e., there are two best models). The similar support for both Boltzmann and Hill models suggests that if we were able to take multiple independent samples in this experimental setup, we would expect to see a considerable amount of variability in the selected best model between samples. For example, for the SOL dataset in P1, the Hill model has a 45 % chance of being the best model, while the Boltzmann model has 36 % chance. The third- and fourth-best models were also supported for some of the given data sets (Δ_AIC_ < 3). However, the Boltzmann or the Hill models were at least three times more likely to be the best model in terms of Kullback–Leibler discrepancy compared to the fourth-best models in all data sets.

**Table 3.**
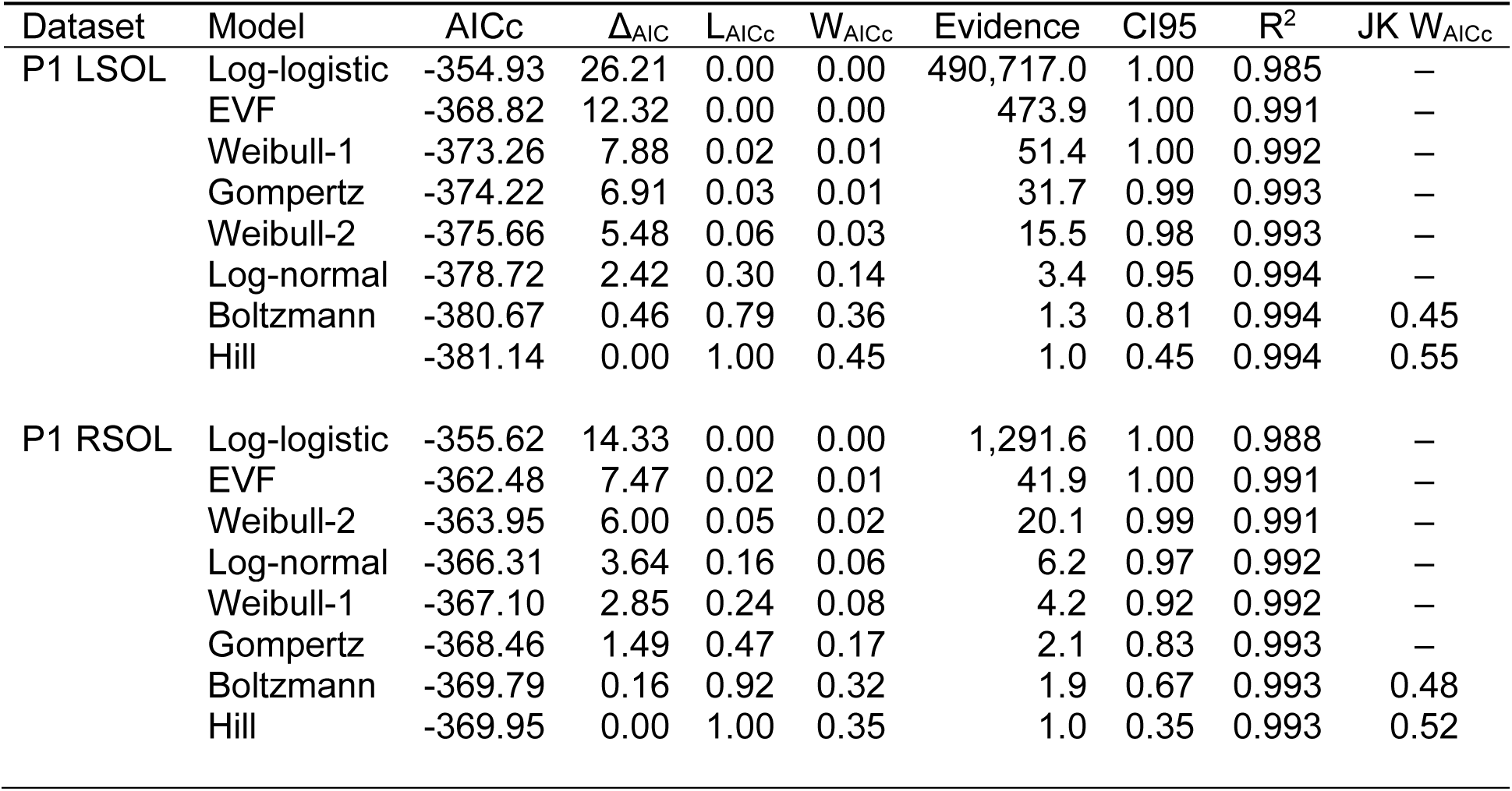
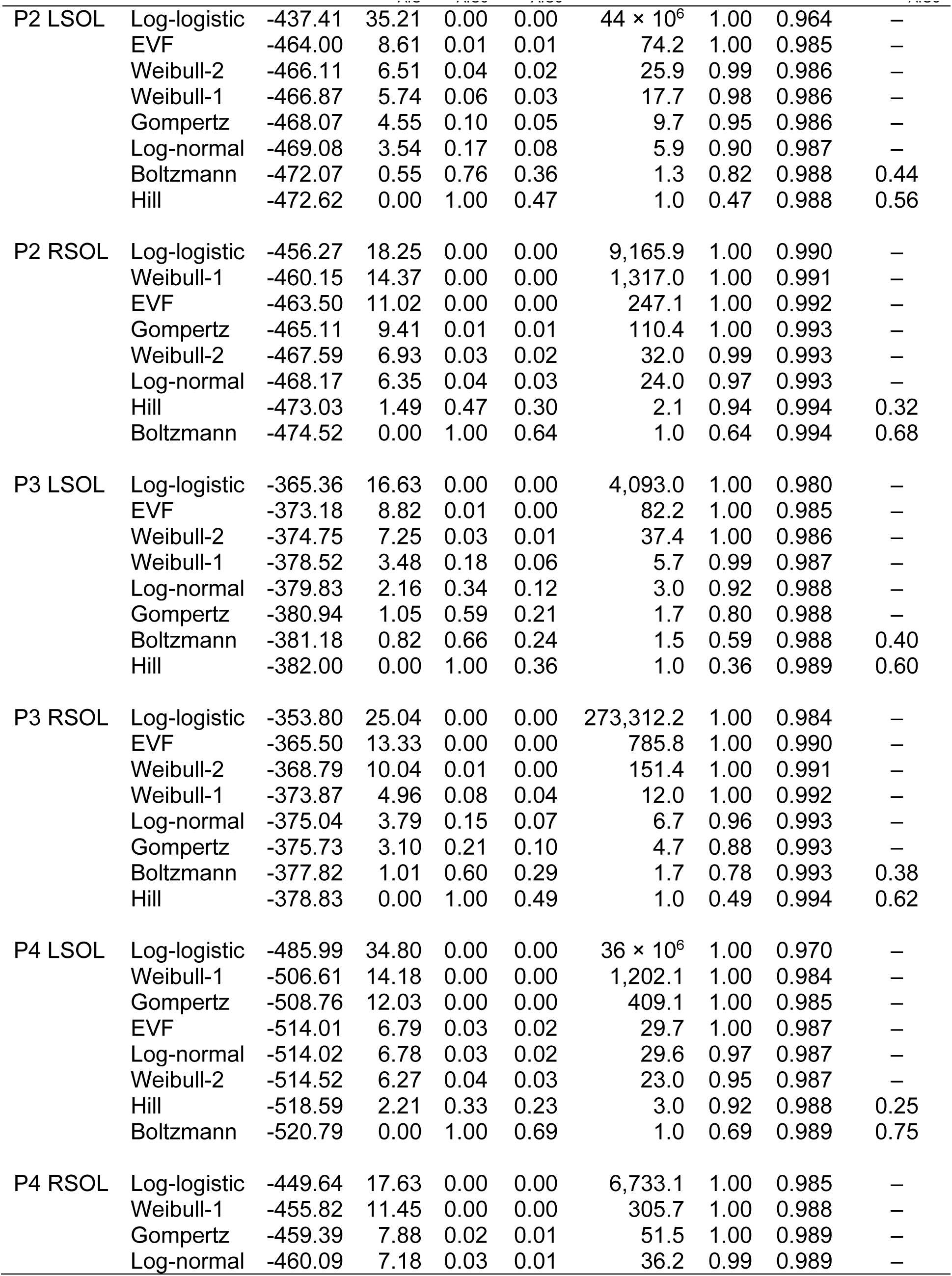
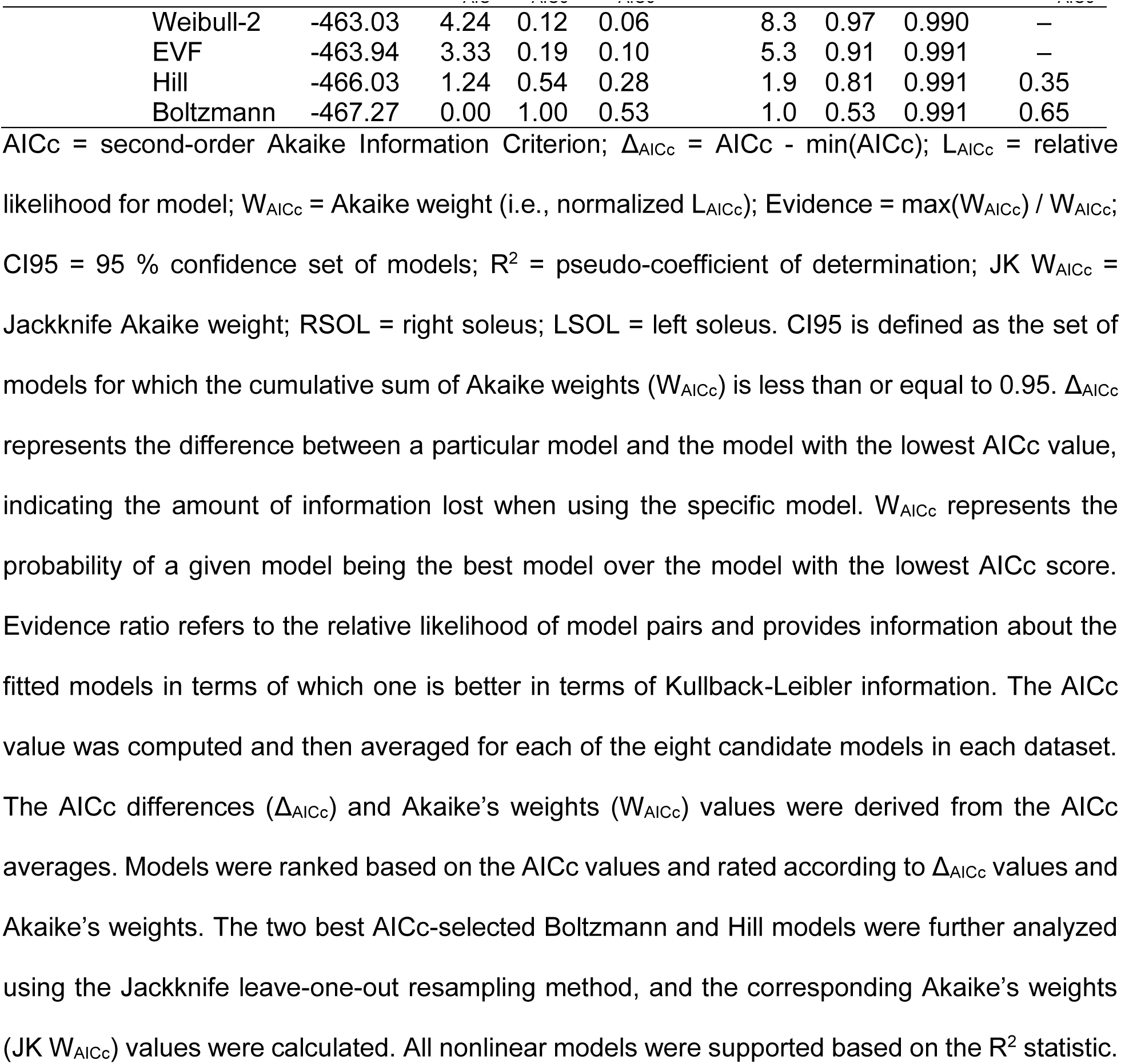
Performance evaluation of models fitted to SOL TEP recruitment curves.

The EVF, Boltzmann, and Hill models were found to be the best to fit the right and left TA TEP input-output recruitment curves for all four protocols (Table 4). The evidence ratio between the two best models ranged from 1.1 to 2.4, while for the best model versus the third-best model ranged from 1.8 to 4.4, suggesting that all these three models are suitable. Although the EVF model has not been fitted before to TEP input-output recruitment curves, we selected this model in our study because the shape of the EVF model is different from Boltzmann and Hill models close to the upper asymptote, but similar at the lower asymptote (Fig 2). The EVF model is suitable to fit TEP input-output recruitment curves with a slower rise of slope reaching a maximum, followed by a more rapid decline. All best models satisfied the goodness-of-fit criterion of coefficient of determination (R^2^ ≥ 0.972) (Tables 3 and 4).

**Table 4.**
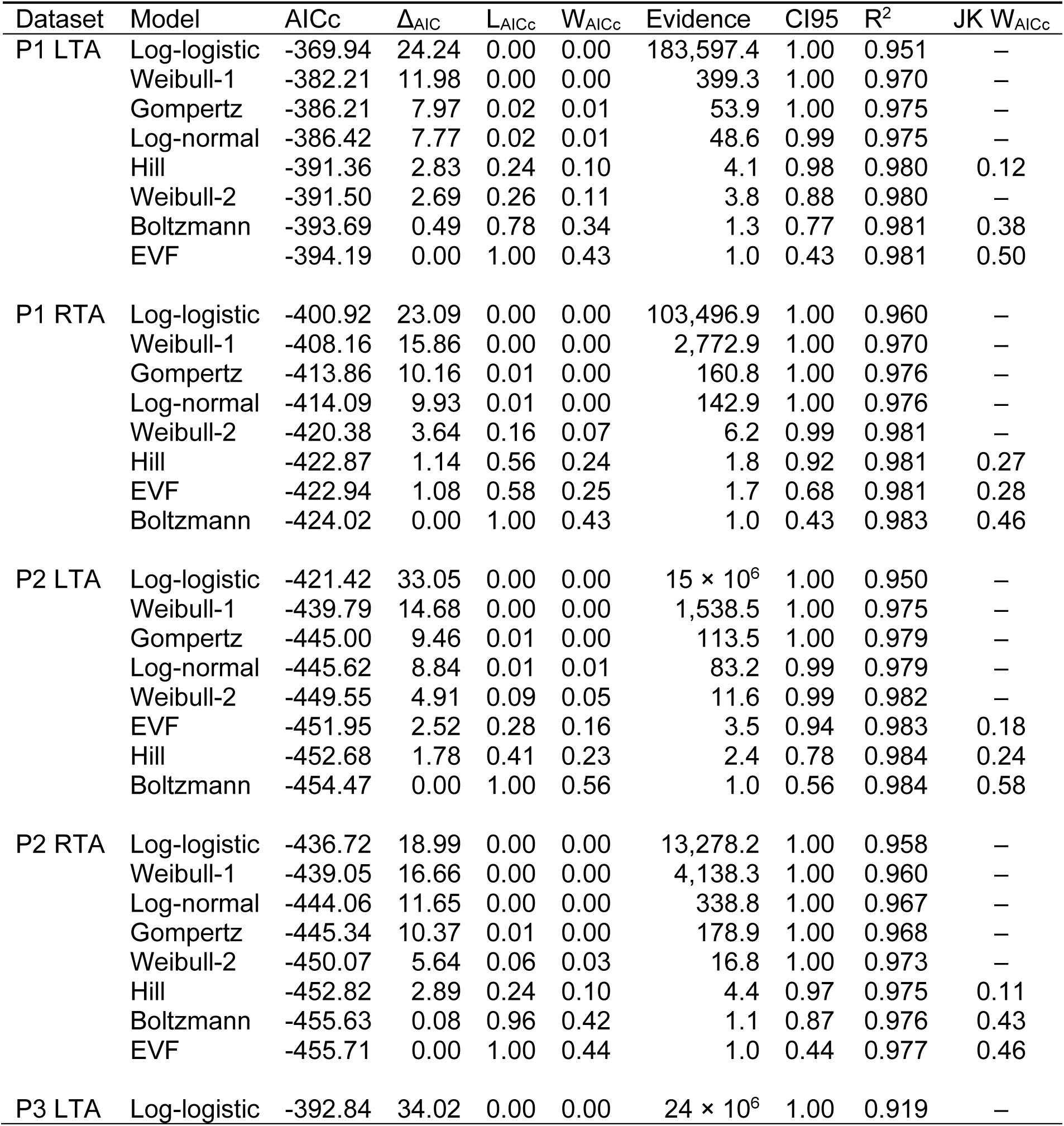
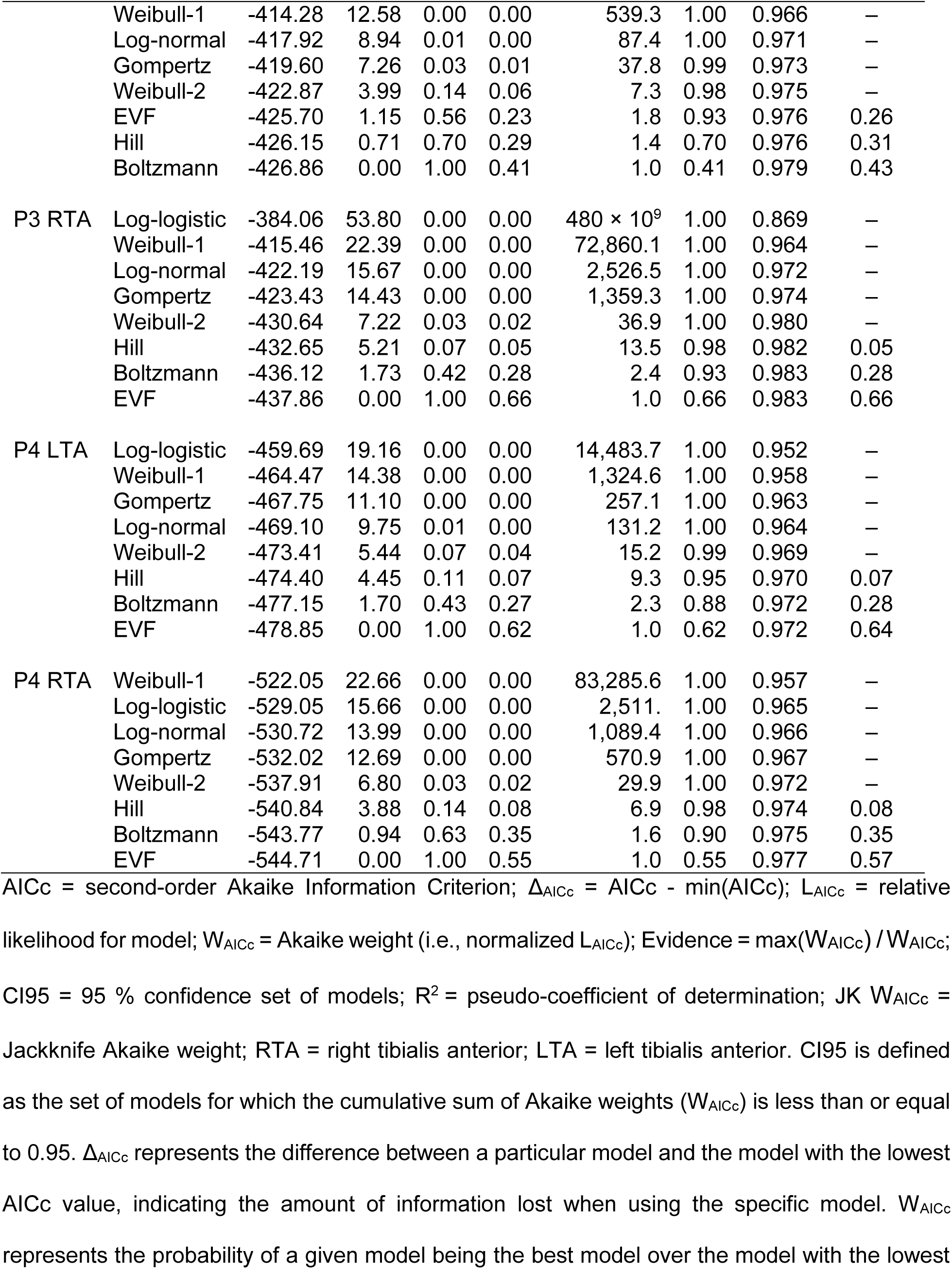
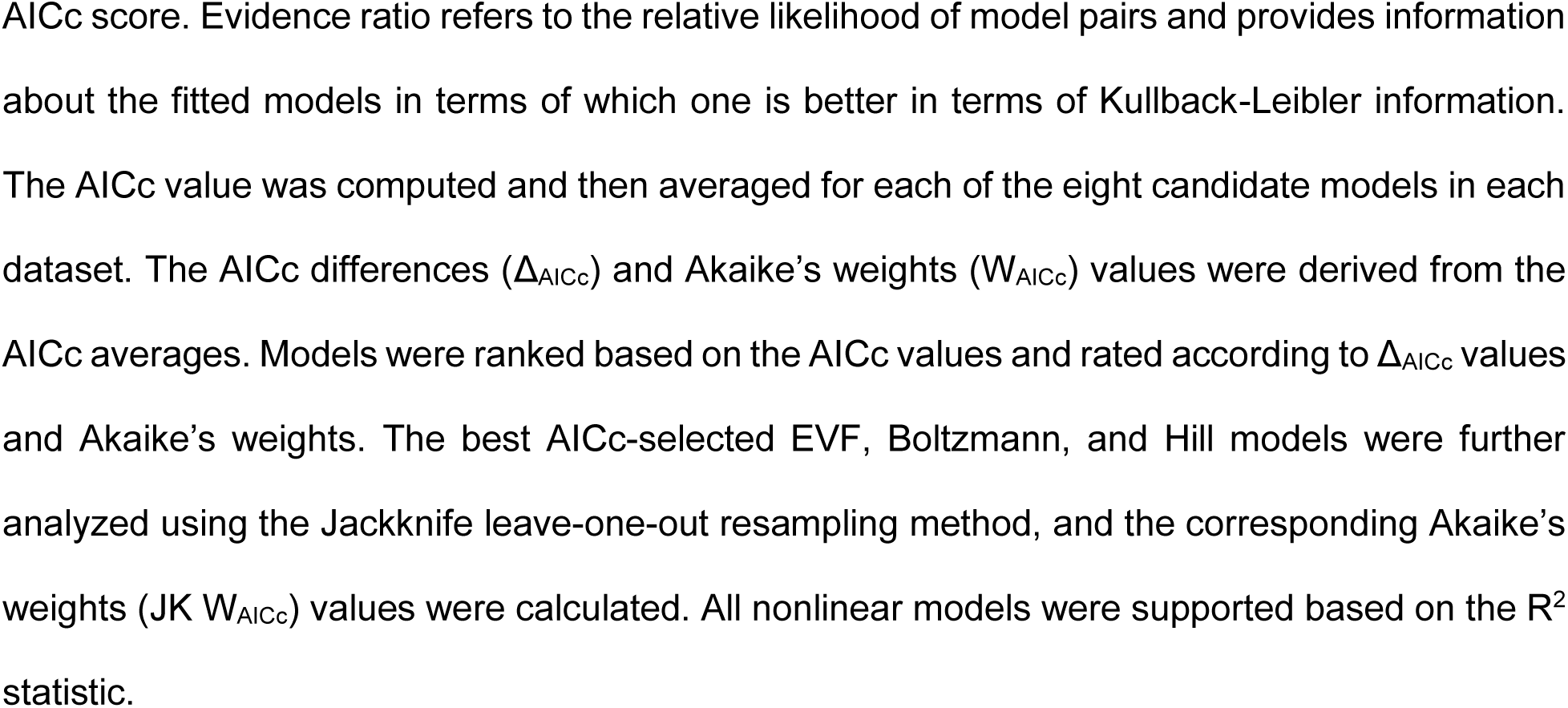
Performance evaluation of models fitted to TA TEP recruitment curves.

The interclass correlation coefficient was ρ = 0.96 for TEPmax, ρ = 0.98 for inflection point and ρ = 1.00 for slope, indicating an excellent agreement between the Boltzmann and Hill models fitted to SOL TEP input-output recruitment curves (Table 5). For TEPmax, 85 % (68 curves out of 80) of the fitted curves had less than 5 % difference, while for slope and inflection point the percentage increased to 95 % and 96 %, respectively (Fig 4). The mean difference (± standard error) between the fitted models was d = 1.56 ± 0.15 % for slope, d = -0.885 ± 0.29 % for inflection point, and d = -3.94 ± 0.78 % for TEPmax. The limits of agreement were smaller for slope (-2.07 – 5.19) compared to those for inflection point (-8.16 to 6.39) and TEPmax (-22.74 to 14.86). For TEPmax, there were some large differences between models, indicating higher values of TEPmax for Hill model compared to that of Boltzmann model. For example, in Fig 5 the fitted curves of the Hill model did not catch the upper limits well mostly in subjects S6 and S9, resulting in higher TEPmax values than that of the Boltzmann model.

**Fig 4.**
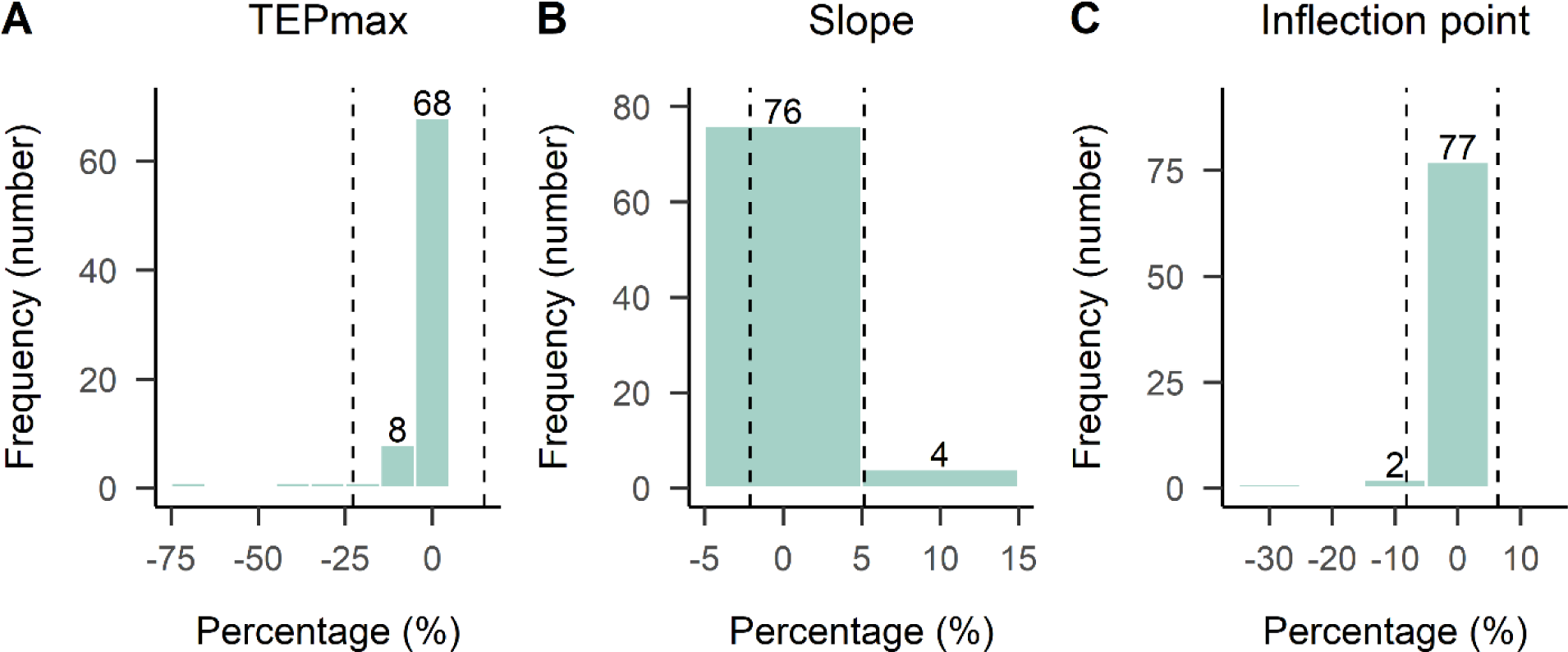
Frequency distribution of percent differences between parameters of best-fit SOL TEP recruitment curve models. TEPmax, slope, and inflection point parameters estimated with the Boltzmann and Hill models fitted to right and left SOL TEP recruitment curves (100 × (Boltzmann - Hill) / (Boltzmann + Hill) / 2). The negative value in the percentage difference indicates that the estimated parameter was larger for Hill model compared to Boltzmann model. Values in abscissa refer to the number of TEP recruitment curves. Vertical dashed lines correspond to limits of agreement band (mean difference ± 1.96 × standard deviation).

**Fig 5.**
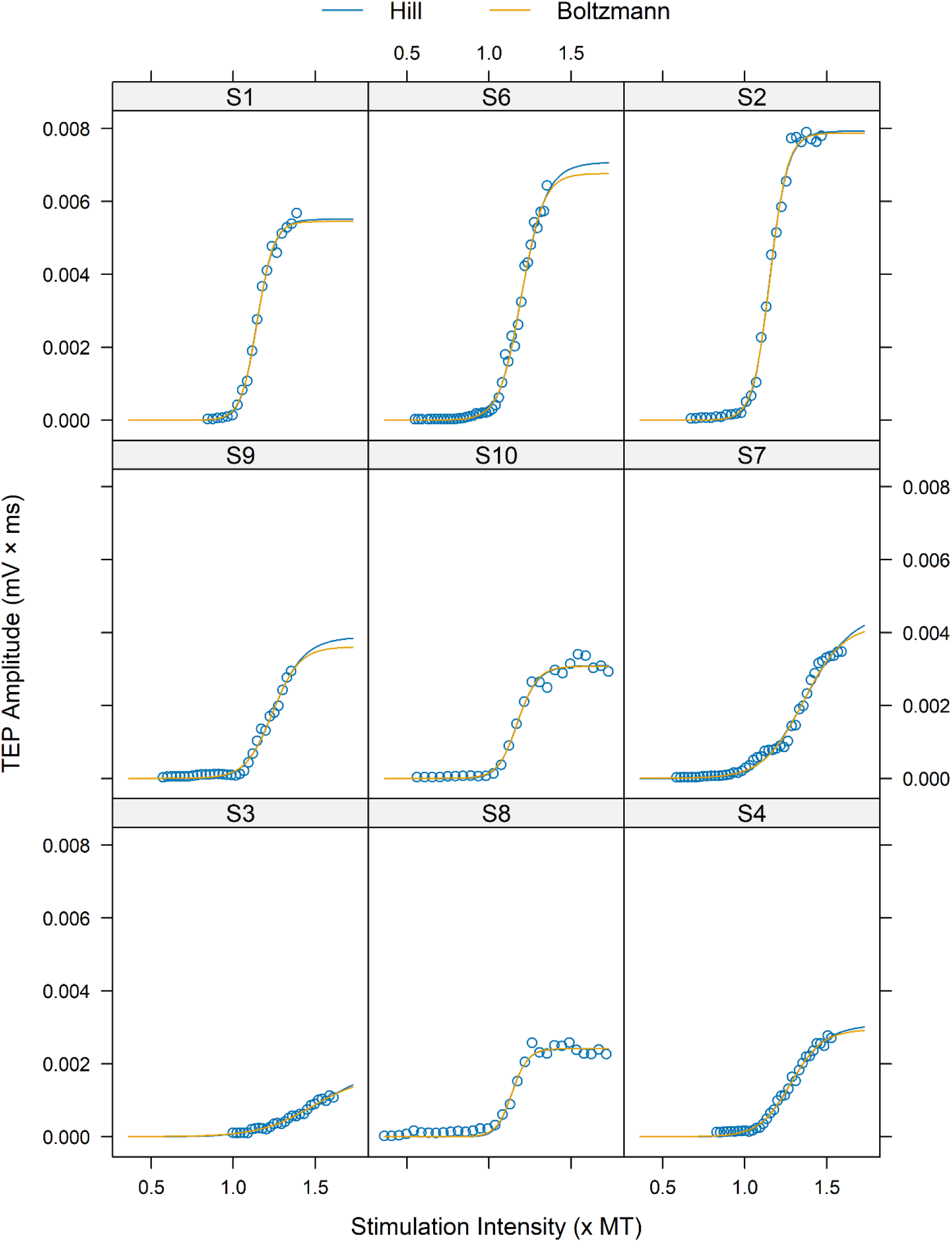
Modelling of SOL TEP recruitment curves using Boltzmann and Hill models. To ensure comparability between curves of different subjects, the units of the X-axis were converted by expressing the stimulation intensity values as multiples of the threshold value (× MT) derived from Boltzmann model. The Hill model did not catch the upper limits well, especially in subjects S6 and S9, resulting in higher TEPmax values than that estimated by the Boltzmann model. The data is for transspinal evoked potentials (TEPs) of the right SOL under protocol P4. For the Boltzmann model, the TEP threshold (B_TH_) is calculated as 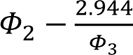, while for the Hill model, the TEP threshold (H_TH_) is calculated as 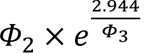.

**Table 5.** Agreement between parameters from best fit TEP recruitment curve models.

| | Parameter | Compared Models | $\rho$ (95 % CI) |
| --- | --- | --- | --- |
| SOL TEP recruitment curves | TEPmax | Boltzmann vs. Hill | 0.96 (0.94 – 0.97) |
|  | Slope | Boltzmann vs. Hill | 1.00 (1.00 – 1.00) |
|  | Inflection point | Boltzmann vs. Hill | 0.98 (0.97 – 0.99) |
| TA TEP recruitment curves | TEPmax | Boltzmann vs. Hill | 0.19 (0.00 – 0.26) |
|  |  | EVF vs. Boltzmann | 0.82 (0.72 – 0.89) |
|  |  | EVF vs. Hill | 0.01 (0.00 – 0.25) |
|  | Slope | Boltzmann vs. Hill | 0.09 (0.00 – 0.33) |
|  |  | EVF vs. Boltzmann | 0.99 (0.98 – 0.99) |
|  |  | EVF vs. Hill | 0.06 (0.00 – 0.30) |
|  | Inflection point | Boltzmann vs. Hill | 0.19 (0.00 – 0.42) |
|  |  | EVF vs. Boltzmann | 0.99 (0.99 – 1.00) |
|  |  | EVF vs. Hill | 0.17 (0.08 – 0.39) |

The interclass correlation coefficient (ρ) for TEPmax, inflection point, and slope parameters estimated by EVF and Boltzmann models fitted to the TA TEP input-output recruitment curves was 0.82, 0.99, and 0.99, respectively (Table 5). Estimates of inflection point, and slope parameters were in excellent agreement between models, with 100% and 95% of the TA TEP recruitment curves, respectively, having less than 15% difference (Fig 6). However, there was a noticeable decrease in agreement when considering TEPmax, with 63 % of the TEP recruitment curves having less than 15% difference (40 curves out of 63). The decreased agreement in TEPmax was reflected in the mean difference values between models, confirming that EVF model differs from Boltzmann close to the upper asymptote. The mean difference (± standard error) between EVF-Boltzmann models was d = -14.0 ± 0.71 % for TEPmax, d = 4.81 ± 0.36 % for slope, and d = 2.0 ± 0.18 % for inflection point. For example, in Fig 7 the fitted curves of the Boltzmann model did not catch the upper limits well in subjects S3, S6 or S5, resulting in higher TEPmax values than that of the EVF model.

**Fig 6.**
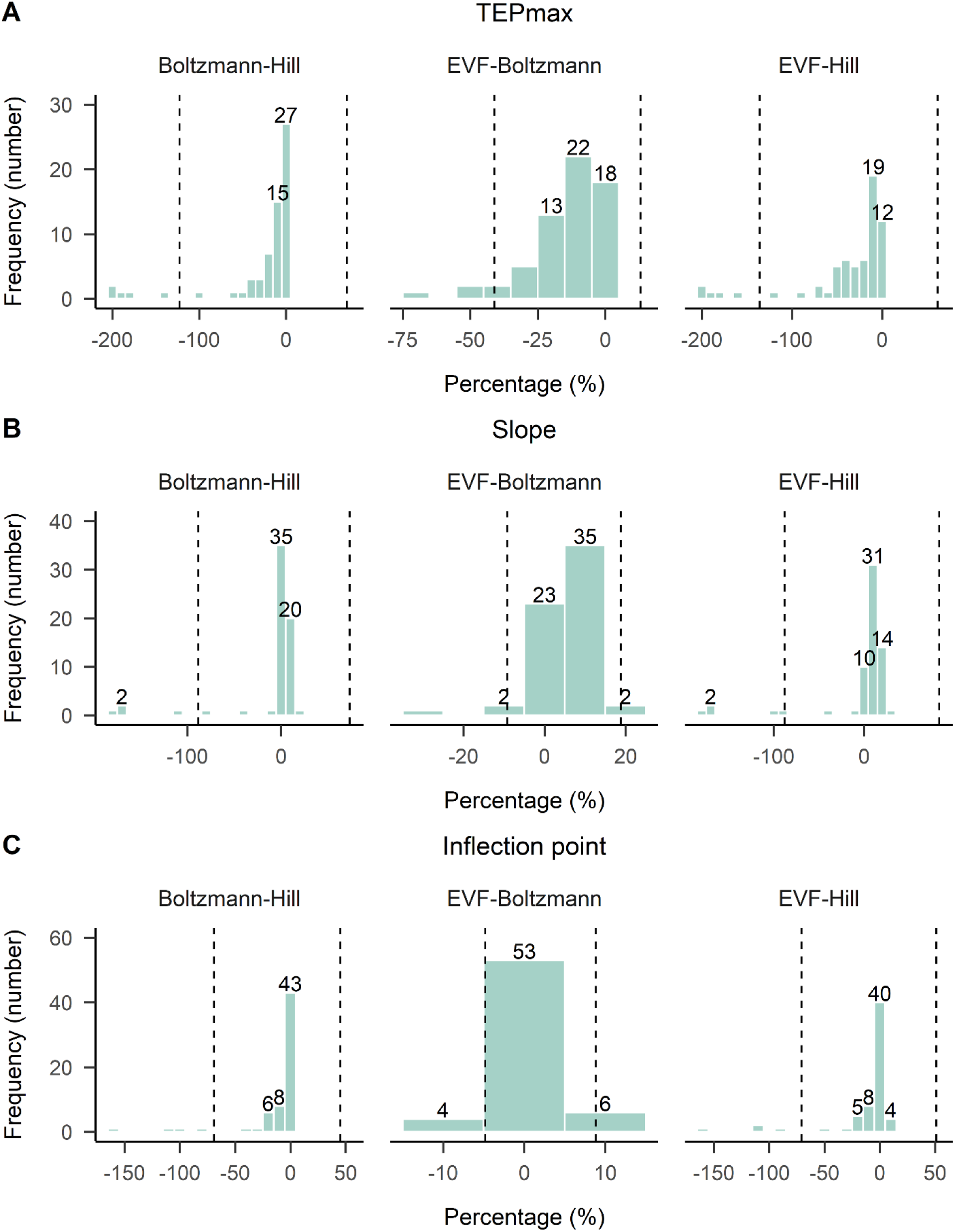
Frequency distribution of percent differences between parameters of best-fit TA TEP recruitment curve models. TEPmax, slope, and inflection point parameters estimated with the EVF, Boltzmann and Hill models fitted to the right and left TA TEP recruitment curves (100 × (model A - model B) / (model A + model B) / 2). The negative value in the percentage difference indicates that the estimated parameter was larger for model B compared to model A (model A – model B). Values in abscissa refer to the number of TEP recruitment curves. Vertical dashed lines correspond to limits of agreement band (mean difference ± 1.96 × standard deviation).

**Fig 7.**
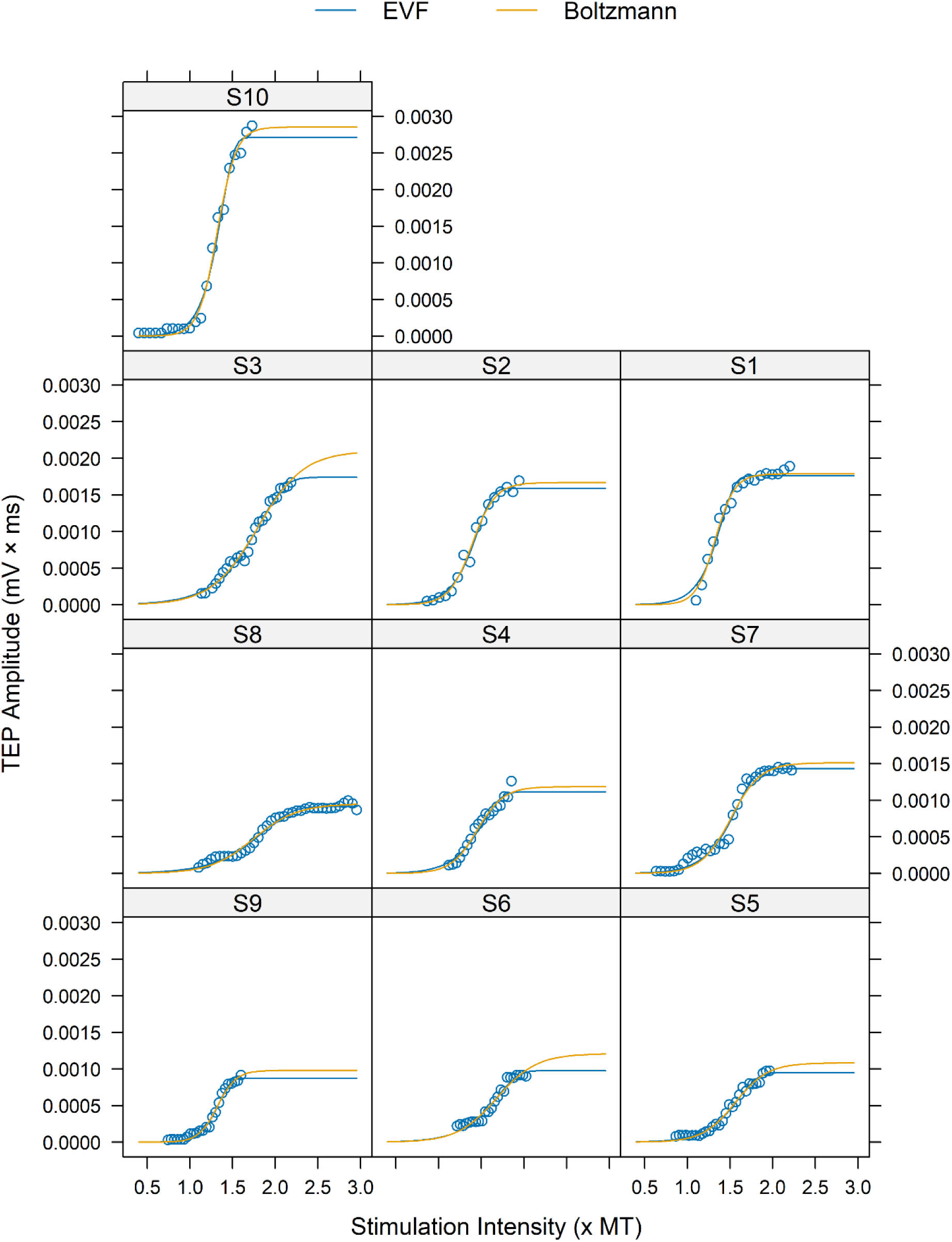
Modelling of TA TEP recruitment curves using EVF and Boltzmann models. To ensure comparability between models, the units of the X-axis were converted by expressing the stimulation intensity values as multiples of the threshold value (× MT) derived from Boltzmann model. The Boltzmann model did not catch the upper limit well, especially in subjects S3 and S6, resulting in a higher TEPmax values than that of the EVF model. The data is for transspinal evoked potentials (TEPs) of the left TA under protocol P1. For the Boltzmann model, the TEP threshold (B_TH_) is calculated as 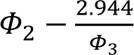, while for the EVF model, the TEP threshold (E_TH_) is calculated as 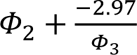

For the Boltzmann-Hill and EVF-Hill models the agreement between estimates of inflection point, slope, and TEPmax parameters was poor, with intraclass correlation coefficients ranging from 0.01 to 0.19 (Table 5). Between Boltzmann and Hill models, 81 % of the curves had less than a 15 % difference in the inflection point, 89 % of the curves had less than a 15 % difference in the slope, and 67 % of the curves had less than a 15 % difference in TEPmax. Likewise, between EVF and Hill models, 82 % of the curves had less than a 15 % difference in inflection point, and 67% of the curves had less than 15 % difference in slope. For TEPmax, only 49 % of the curves exhibited less than 15 % difference (31 curves out of 63). Between Boltzmann and Hill models the mean difference was d = -26.4 ± 2.51% for TEPmax, d = -7.6 ± 2.11 % for slope, and d = -12.0 ± 1.49 % for inflection point, and between EVF and Hill models the mean difference was d = -37.0 ± 2.57 % for TEPmax, d = -2.21 ± 2.23 % for slope, and d = -9.8 ± 1.59 % for inflection point. The limits of agreement were smaller for EVF-Boltzmann models (slope: -9.19 – 18.81; inflection point: -4.82 – 8.82; TEPmax: -41 – 13) compared to EVF-Hill (slope: -87.91 – 83.49; inflection point: -70.7 – 51.1; TEPmax: -135.6 – 61.6) and Boltzmann-Hill models (slope: -88.4 – 73.2; inflection point: -69.1 – 45.1; TEPmax: -122.6 – 69.8).

### Models’ assumptions

The Shapiro-Wilks test was performed to evaluate whether the residuals of the best models followed a normal distribution. The results showed that among the best models for the SOL TEP input-output curves, the null hypothesis of normal distribution of residuals was rejected in 29 and 27 cases (out of 80) for the Boltzmann and Hill models, respectively (p < 0.05). Similarly, for the TA TEP input-output recruitment curves the null hypothesis of residual normality was rejected in 14, 16, and 11 cases (out of 65) for the Boltzmann, Hill, and EVF models, respectively (p < 0.05). A jackknife leave-one-out method was applied to the best models to confirm the presence of influential data points. The jackknife procedure is a resampling technique that systematically removes each observation and evaluates the impact on parameter estimation. We decided not to exclude any of the influential data as an outlier, because each data point represented an average of four replicates. The jackknife Akaike weights (JK W_AICc_) confirmed the uncertainty to select only one best mathematical model to fit the TEP recruitment curve (Tables 3 and 4). The Ljung-Box test was used to test the null hypothesis of no residual autocorrelation against the alternative hypothesis of non-zero coefficient in a first-order autoregressive correlated errors model. The results showed that among the best models for SOL TEP input-output curves, the null hypothesis was rejected in 40 cases for the Boltzmann model, and in 38 cases for the Hill model (out of a total of 80 cases) (p < 0.05). Similarly, for the best models for the TA TEP input-output curves, the null hypothesis of no residual autocorrelation was rejected in 45 cases for the Boltzmann model, in 44 cases for the Hill model, and in 46 cases for the EVF model (out of a total of 63 cases) (p < 0.05).

## Discussion

In this study, we found for the first time that the distribution of the recruitment gain is muscle dependent based on the best fit sigmoidal model. The SOL TEP recruitment input-output curve was best fitted with the symmetric Boltzmann model, while the asymmetric EVF model was more suitable for the TA TEP recruitment input-output curve. The selection of an appropriate sigmoidal model is important in neurophysiological research because the predicted parameters derived from the input-output curves are biomarkers for understanding the physiological or pathological behavior of neuronal networks and post-treatment recovery [7,21,48–51].

### Physiological applications of the findings

As evident from the results, the form of the TEP input-output relation is sigmoidal. A sigmoidal TEP input-output curve suggests that transspinal stimulation of increasing strength activates transynaptically motoneurons in an orderly manner, from low-threshold to high-threshold, with increasing TEP amplitudes from threshold to saturation [52–54]. However, the maximal TEP value at saturation cannot be regarded as the maximal response to pure excitatory neuronal events but rather a net motor output of the spinal cord representing the difference between net excitation and net inhibition acting on motoneurons and interneurons brought up by transspinal stimulation [28,53,55]. The overall effect of the heterogeneous distribution of the net input within the pool, the heterogeneity of motoneuron population and neural structures, and the heterogeneous distribution of their properties within the pool caused the sigmoidal input-output relation [13,56–58]. The same monotonic and nonlinear form of input-output relation (sigmoidal) was found for SOL and TA TEPs. However, a symmetrical bell-shaped curve of the recruitment gain was found only for SOL TEPs, while the recruitment gain of the TA TEPs was negatively skewed. This may be related to a different proportion of slow vs. fast motoneurons being recruited in the SOL and TA TEPs leading probably to a skewed distribution of neuronal excitability spectrum across the motoneuron pool [48,59].

The Boltzmann model fitted to TEP input-output curve explicitly implies a symmetrically distributed neuronal excitability spectrum across motoneuron pool [28,55], where the greatest change in the amount of net motor output (recruitment gain) is confined to occur at 50 % of the TEPmax – i.e., smaller increments of transspinal stimulation strength is needed to increase net motor output [26,28]. However, a skewed distribution of the relative excitabilities of motoneurons within the pool is possible [22,23,48]. Our results showed that a symmetric recruitment gain does not hold for TA TEP input-output recruitment curves. The TA TEP input-output recruitment curves are best fitted with the asymmetric EVF model. The EVF model describes an asymmetric recruitment curve where the maximum recruitment gain is confined to occur at 63.2 % of the TEPmax, but otherwise is similar to the Boltzmann model.

### Benefits and pitfalls in empirical modeling TEP input-output recruitment curve in neurophysiology research

The Boltzmann and Hill models have been the models of choice for TEP input-output recruitment curves, regardless of the target muscle. Both models yield nearly identical fits when the stimulation-response dataset adequately represents the full range of the recruitment curve – i.e., when the lower and upper limits of the input-output relationship are well-defined. Moreover, the parameters’ interpretation is consistent between models (TEPmax, S50, slope). Inaccuracies in parameter estimation can arise when fitting a stimulation-response dataset with imperfect sampling using either a Hill or Boltzmann model. For instance, factors such as subject discomfort or limitation in the maximum stimulator output may lead to scenarios during which responses are not recorded at supramaximal intensities. This may result in an insufficient number of data points at the upper limit of the recruitment curve. The results showed that the Hill model was more susceptible to providing inaccurate parameter estimates compared to the Boltzmann model when the upper limit of the SOL TEP input-output recruitment curve is poorly determined by the experimental data. Therefore, it is recommended to use the Boltzmann model to fit the SOL TEP input-output recruitment curves. In cases where convergence problems arise with the Boltzmann model, the Hill model can be used, provided the stimulation-response dataset is well-defined [24]. High-order polynomial and piecewise polynomial models have also been fitted to SOL or TA TEP recruitment curves. Examples include sixth-order polynomials [22,23,60] or quintic splines within the generalized additive model framework [61]. Compared with the proposed three-parameter nonlinear models, a competing high-order polynomial model lacks parsimony and interpretability. Using a sixth-order polynomial to describe the same data would have the disadvantage of requiring more than three parameters, none of which would have a natural physical interpretation. As a consequence, additional computations would have needed to derive the neurophysiologically important parameters for describing the TEP recruitment curves (slope, inflection point, TEPmax, TEP threshold), which include computing the first and the second derivatives of the fitted polynomial and finding local minima and maxima [22,23,60,61].

Another disadvantage of fitting high-order polynomials is their interpolation behavior, which can yield unrealistic responses. For instance, when a competing sixth-order polynomial model is fitted to a TEP input-output recruitment curve, the model’s values can become negative, lacking any biologically meaningful interpretation (for example see Fig. 1 in [22]). To mitigate such unrealistic behavior, a six-order polynomial model is sometimes used to fit only the ascending limb of the TEP input-output recruitment curve, representing an approximately linear increase in TEP amplitude with increasing stimulus intensity. However, the decision regarding which data to include depends on the methods used to estimate the threshold intensity. This estimation often involves ad hoc statistical procedures such as surpassing the average plus three times the standard deviation of the baseline [23]. Therefore, polynomials are only occasionally useful to model TEP recruitment curves and were not part of the proposed models in this study.

A limitation of the study was that the nonlinear models were fitted to sample means, and the variability of TEP replicates at each stimulation intensity was not taken into consideration. In future work, it may be useful to fit the nonlinear models to the stimulus-response data using a mixed-effects modelling approach that allows estimation of the average behavior of multiple TEP recruitment curves [40]. Additionally, the overall agreement between EVF and Boltzmann models, and between Boltzmann and Hill models for TA and SOL TEP recruitment curves, respectively, suggests that in research studies, more than one model can be used based on the muscle from which responses are recorded. Future research could seek to address this issue by conducting model averaging based on AIC values. AIC-based model averaging combines information from multiple models, weighing them based on their AIC values [30,31].

## Conclusions

In this study, an empirical comparison between eight sigmoidal models to fit the TEP recruitment curve was conducted. Experimental studies aiming to predict parameters of interest from TEP recruitment curves should consider comparing different sigmoidal functions based on information-theoretic criteria. We suggest using the Boltzmann model for fitting the SOL TEP recruitment curve and the EVF model for fitting the TA TEP recruitment curve.

## Data availability

All data files are available from the corresponding author upon request.

## Author Contributions

Andreas Skiadopoulos, Maria Knikou: Conceptualization, Writing – original draft, reviewing & editing.

Andreas Skiadopoulos: Methodology, data analysis, statistics. Maria Knikou: Funding acquisition, supervision.

